# CRISPR/Cas9-mediated resistance to cauliflower mosaic virus

**DOI:** 10.1101/191809

**Authors:** Haijie Liu, Cara L. Soyars, Jianhui Li, Qili Fei, Guijuan He, Brenda A. Peterson, Blake C. Meyers, Zachary L. Nimchuk, Xiaofeng Wang

## Abstract

Viral diseases are a leading cause of worldwide yield losses in crop production. Breeding of resistance genes (*R* gene) into elite crop cultivars has been the standard and most cost-effective practice. However, *R* gene-mediated resistance is limited by the available *R* genes within genetic resources and in many cases, by strain specificity. Therefore, it is important to generate new and broad-spectrum antiviral strategies. The CRISPR-Cas9 (clustered regularly interspaced palindromic repeat, CRISPR-associated) editing system has been employed to confer resistance to human viruses and several plant single-stranded DNA geminiviruses, pointing out the possible application of the CRISPR-Cas9 system for virus control. Here we demonstrate that strong viral resistance to cauliflower mosaic virus (CaMV), a pararetrovirus with a double-stranded DNA genome, can be achieved through Cas9-mediated multiplex targeting of the viral coat protein sequence. We further show that small interfering RNAs (siRNA) are produced and mostly map to the 3' end of guide RNAs (gRNA), although very low levels of siRNAs map to the spacer region as well. However, these siRNAs are not responsible for the inhibited CaMV infection because there is no resistance if Cas9 is not present. We have also observed edited viruses in systematically infected leaves in some transgenic plants, with short deletions or insertions consistent with Cas9-induced DNA breaks at the gRNA target sites in coat protein coding sequence. These edited coat proteins, in most cases, led to earlier translation stop and thus, non-functional coat proteins. We also recovered wild-type CP sequence in these infected transgenic plants, suggesting these edited viral genomes were packaged by wild-type coat proteins. Our data demonstrate that the CRISPR-Cas9 system can be used for virus control against plant pararetroviruses with further modifications.

## Introduction

Viral resistance in plants is mediated by multiple mechanisms consisting of effector-triggered immunity (ETI), loss or mutation of host genes essential for viral infection (Kang et al., 2005; Maule et al., 2007; Wang and Krishnaswamy, 2012), and RNA interference (RNAi)-based innate immunity targeting viral RNAs (Ding, 2010), among others. ETI is conferred through the recognition of viral encoded elicitors by dominant *R* gene products (Kang et al., 2005; Maule et al., 2007; Wang and Krishnaswamy, 2012). The best-studied ETI-based viral resistance is *N* gene-mediated resistance to tobacco mosaic virus. *N* gene encodes a protein with typical R protein features: a Toll, interleukin-1 related region (TIR), a nucleotide binding site (NBS) domain, and a leucine-rich repeat (LRR) domain. The *N* gene product recognizes the helicase-domain of the replication protein to trigger the hypersensitive response (Erickson et al., 1999; Padgett et al., 1997). Among 14 identified recessive *R* genes, 12 encode mutants of either eukaryotic initiation factor 4E (eIF4E) or its isoform eIF(iso)4E (Wang and Krishnaswamy, 2012). Both eIF4E and eIF(iso)4E are involved in translation and function redundantly in plant growth and development. However, one of them but not both is required for infection of some viruses (Wang and Krishnaswamy, 2012). At least for potyviruses, a physical interaction between wild-type (wt) eIF(iso)4E or eIF4E and a specific viral protein, termed viral protein, genome-linked (VPg), has been demonstrated. *R* gene-encoded eIF4E or eIF(iso)4E mutants fail to interact with VPg and conversely, resistance-breaking potyvirus isolates bear mutations in VPg (Wang and Krishnaswamy, 2012). In crop plants, ETI may be highly specific to certain strains or may not be durable due to viral mutation in R-protein recognized elicitors. Recessive *R* genes are durable, however, not many are available. In addition, viral genomes often encode suppressors of host RNAi machinery (termed viral suppressors of RNA silencing) rendering RNAi-based resistance ineffective (Anandalakshmi et al., 1998; Incarbone and Dunoyer, 2013; Kasschau and Carrington, 1998) and in many cases, allowing the co-infecting viruses to replicate to much higher levels that lead to detrimental diseases in host plants (Anandalakshmi et al., 1998).

The CRISPR-Cas (Clustered, regularly interspaced short palindromic repeats-CRISPR-associated) system is an adaptive immune defense mechanism employed by bacteria and archaea to fight against invading viruses and foreign nucleic acid materials (Makarova et al., 2011; Sander and Joung, 2014). The engineered CRISPR-Cas system includes a guide RNA (gRNA) and a Cas nuclease (Jinek et al., 2012). The gRNA, via the 20-nucleotide spacer region, anneals to the complementary strand of the targeted double-stranded DNA (dsDNA) and recruits the Cas nuclease to make dsDNA breaks at the target site. Mutational insertions and/or deletions are introduced when the breaks are joined and repaired incorrectly. The class II CRISPR-Cas9 system has been engineered to confer resistance to various human viruses (Price et al., 2016) and plant geminiviruses (Ali et al., 2015; Baltes et al., 2015; Chaparro-Garcia et al., 2015; Ji et al., 2015). In the majority of these cases, a single gRNA is used and expressed at high levels. It has been reported that HIV mutants with mutations at the gRNA targeting sites escape the CRISPR-Cas9-mediated viral resistance, suggesting multiple gRNAs could be a better choice for virus control because they are harder to overcome by viruses (Wang et al., 2016b; Wang et al., 2016c). Given that gRNAs are highly structured RNAs with portions of dsRNA, it is possible that gRNAs can be targeted by cellular RNAi machinery in transgenic plants or animals and thus, the abundance of gRNAs could be under the regulation of cellular RNAi. However, it is unknown whether small interfering RNAs (siRNA) are generated in eukaryotic cells expressing prokaryotic gRNAs.

Cauliflower mosaic virus (CaMV) is a plant pararetrovirus and has a dsDNA genome with three breaks, which are repaired in the nucleus prior to transcription during viral infection. CaMV mainly infects plant species in the *Brassicaceae* family, including *Arabidopsis thaliana*, and several *Nicotiana* species (Cecchini et al., 1998; Schoelz et al., 1986). We report here that expressing multiple gRNAs targeting the CaMV coat protein (CP) coding sequence confers resistance to the virus in Arabidopsis transgenic plants. We have identified various large deletions of *CP* fragments as early as three days post infection (DPI). However, we also show that edited viruses can escape the infection sites to move systematically in some transgenic plants. We have also found that small RNAs ranging from 21-24 nucleotides (nt) are generated from gRNAs. The majority of siRNAs are mapped to the 3’ end of the gRNA backbone region and very infrequently to the spacer region, which anneals to the CaMV *CP* coding sequence. Our data demonstrate that the CRISPR-Cas9 system can be used for virus control against plant pararetroviruses.

## Results and Discussion

To test the applicability of the CRISPR-Cas9 system in achieving resistance to CaMV, we designed six individual gRNAs that targeted distinct sites in the *CP* coding sequence (Fig. S1 and Table 1) (Chapdelaine and Hohn, 1998). Each gRNA contained 20 nucleotides that are either identical or complementary to the *CP* sequence, followed by a proto-spacer adjacent motif (PAM) sequence (Fig. S1 and Table 1). Each gRNA was expressed from its own Pol III *U6* promoter (Nekrasov et al., 2013). The set of six gRNAs was cloned as a linear array into the *pCUT3* binary vector to create the *pCUT CP* construct, in which the Arabidopsis *UBQ10* promoter is used to express *Cas9* (Peterson et al., 2016). In addition, we made a *pCUT CP-No Cas9* construct, in which the *Cas9* coding sequence was removed from *pCUT CP*. This construct serves as a negative control to demonstrate that the presence of CaMV-targeting gRNAs alone is unable to affect CaMV infection. Each of the two constructs was transformed into wild-type (wt) Arabidopsis Col-0 ecotype plants to make *CUT CP* or *CUT CP/No Cas* transgenic plants. It should be noted that the transgenic plants harboring the above constructs grew normally and were fully fertile, indicating that the *CP* gRNAs or Cas9 and gRNA combinations did not negatively impact plant growth. As a specificity control we also included a transgenic line *CUT GLV* (Peterson et al., 2016), which contains a 14 gRNA unit array targeting seven members of the Arabidopsis *GOLVEN* (*GLV*) gene family (Fernandez et al., 2013).

**Table 1.**
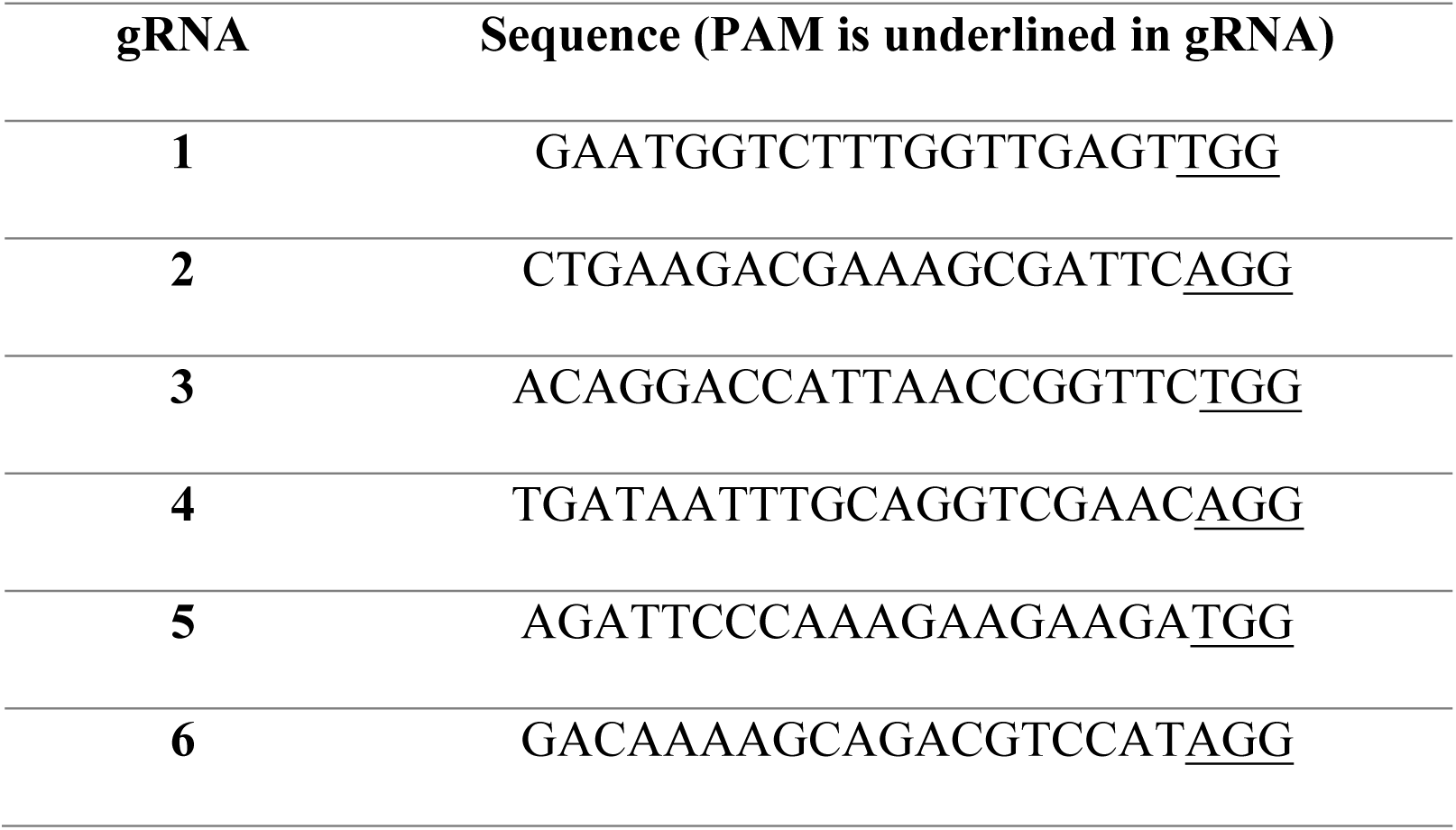
Cas9 target sites selected in the CaMV *CP* gene.

We infected transgenic and Col-0 plants with partially purified CaMV virions by mechanical inoculation and monitored symptom progression in systemic leaves. In Col-0, *CUT GLV*, and *CUT CP/No Cas9* plants, vein clearing and chlorosis symptoms appeared on the inoculated leaves around 10 DPI. By 14 DPI, chlorosis extended to the majority of emerging systemic leaves. By three weeks, infected plants were highly chlorotic and stunted in 19, 16, and 17 of 20 inoculated Col-0, *CUT GLV*, and *CUT CP/No Cas9* plants, respectively (Fig. 1A). This indicated that neither Cas9 nor gRNAs targeting non-viral genomes confer resistance. Conversely, when 20 plants of two transgenic lines, *CUT CP*-1 and -2, were tested, 17 and 18 plants remained symptomless at 20 DPI (Fig. 1A). Similar results were obtained in repeated infection assays where 81% (*CUT CP*-1, Fig. S2A) and 96% (*CUT CP*-2, Fig. S2A) plants remained symptomless. CaMV was readily detectable in systemic leaves from infected WT, *CUT GLV*, and *CUT CP/No Cas9* lines by real time PCR for the *CP* gene (Fig. 1B) or by PCR for viral genes encoding CP or reverse transcriptase (*RT*, ORFV) (Fig S2B), and in the *CUT CP* lines that displayed symptoms. In contrast, viral DNA was undetectable in symptomless *CUT CP* lines (Fig. 1B, and Fig. S2B). The number of symptomless *CUT CP/No Cas9* control plants was similar to that of CaMV-infected wt and *CUT GLV*, suggesting that gRNAs complementary to invading viruses without Cas9 do not induce resistance (Fig. 1A).

**Figure 1.**
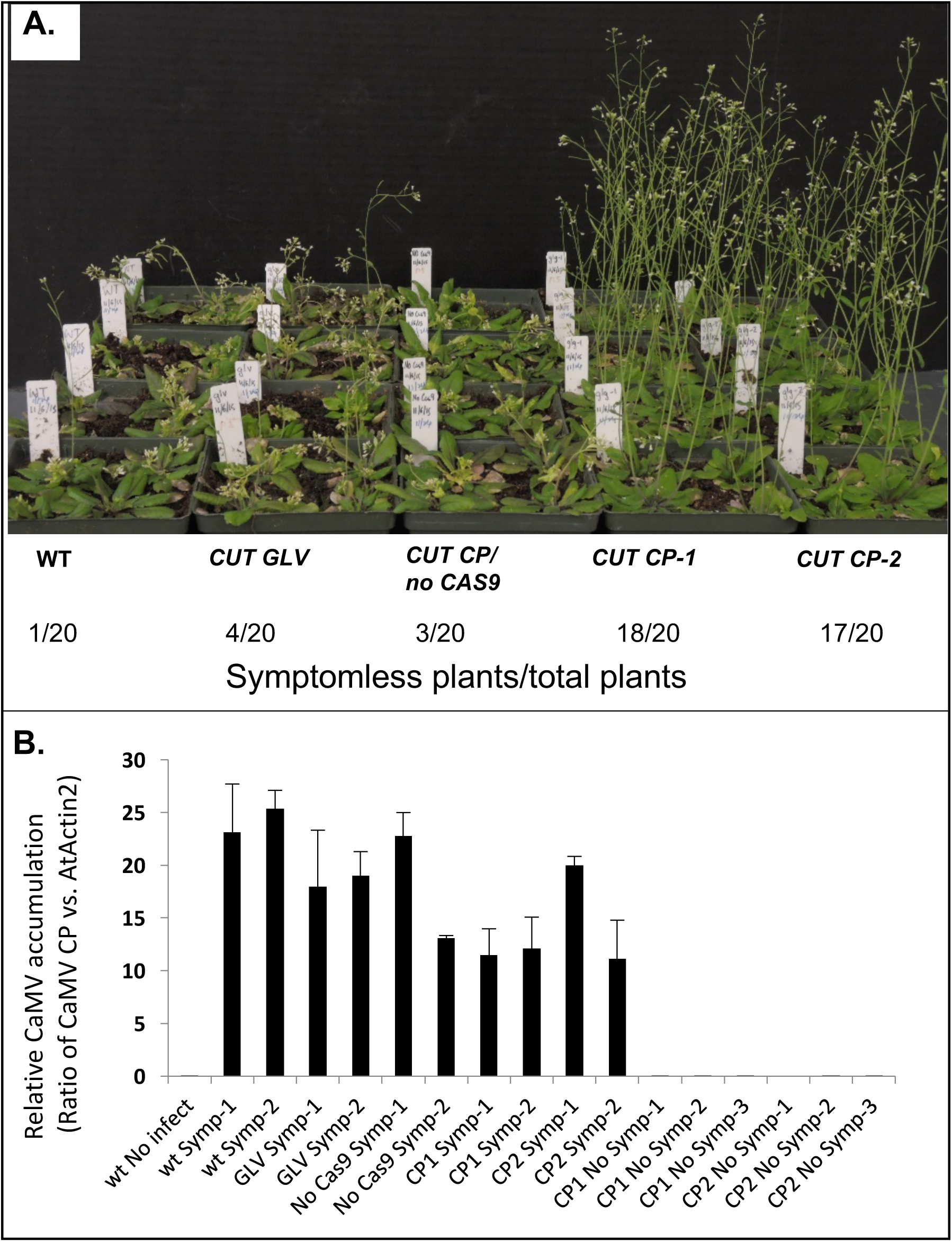
CRISPR-Cas9-based CaMV resistance. A. Cas9 is required for CRISPR-Cas9 system-mediated resistance. From left to right in rows, WT, *CUT GLV*, *CUT CP/No Cas9, CUT CP-1*, *CUT CP-2*. Photos were taken at 20 DPI. Numbers below summarize the number of plants without symptoms in all inoculated plants. B. Quantification of CaMV levels during Cas9-mediated resistance. The ratio of *CP* against an internal *AtACTIN2* standard was calculated to determine the relative accumulation of CaMV. Error bars correspond to means ± SD of two independent experiments with three replicates per sample. Symptomatic plants of wt, *CUT GLV*, CUT NoCas9, CUT CP1, CUT CP2

During CaMV infection, 35S and 19S RNAs are produced in the nucleus and subsequently translocated to the cytoplasm for translation. The 35S RNA also serves as a template to produce progeny dsDNAs in the cytoplasm. Although a remote possibility, viral RNAs might be targeted by small interfering RNAs (siRNAs) generated from gRNAs and if so, virus resistance could possibly be due to RNAi and not the CRISPR-Cas9 system. To test this notion, we performed small RNA deep sequencing on CaMV-inoculated or mock-inoculated leaves at three DPI in wt, *CUT GLV*, *CUT CP* plants. In particular, we focused on small RNAs that are 21, 22 and 24 nucleotides (nt) in length, because they are derived from the activity of different DICER-LIKE (DCL) proteins and are involved in RNAi (Seo et al., 2013). We considered these sizes of small RNAs as siRNAs in this report. In contrast, other sizes of small RNAs are more likely to be degradation products of longer RNAs. In the absence of infection, plants expressing the *CP* gRNA arrays accumulated three siRNAs in one *CUT CP* transgenic line, *CUT CP*-1 (Fig. 2A, the panel of *pCUT CP*-1 uninfected). The copy number of each siRNAs was very low (1-10 copies among 10 million small RNA reads). Arabidopsis leaf tissues that were inoculated with CaMV showed abundant siRNA production from viral RNAs at three days post infection (DPI). The highest abundant siRNAs were located at the 3’ end of CaMV genome (Fig. 2A). Compared to wt and *CUT GLV* plants, siRNAs mapped to the CaMV DNA sequence did not show higher abundances in *CUT CP* leaves inoculated with CaMV (Fig. 2A). Therefore, the siRNA-mediated antiviral effect was not triggered by the introduction of multiple gRNAs in Arabidopsis transgenic plants.

**Figure 2.**
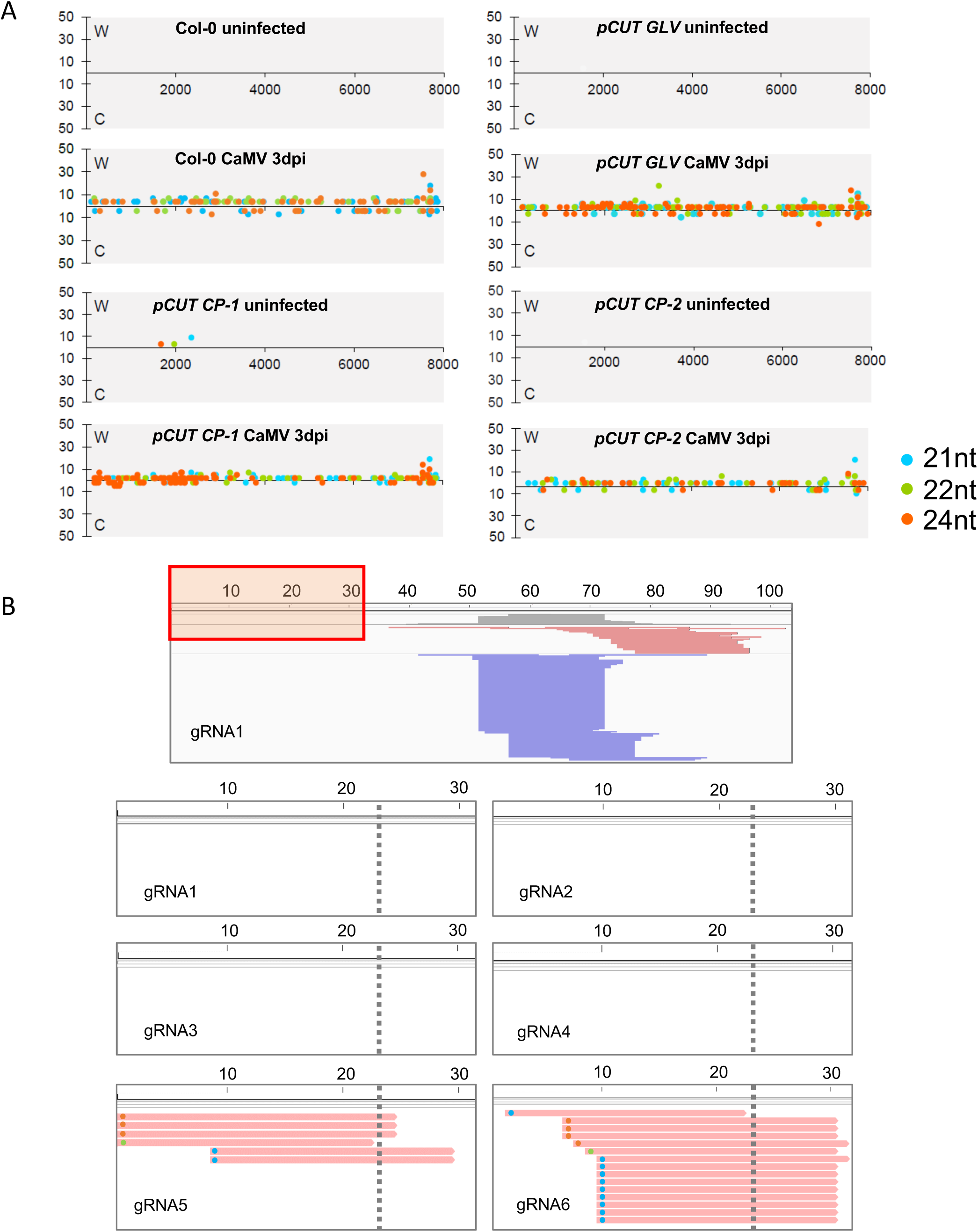
Small RNAs generated from CaMV RNA and gRNAs. A. Small RNAs that are mapped to the CaMV genome. Small RNAs of different sizes are indicated by different colored dots, displaying only sizes potentially produced by Dicer (21/22/24 nt). The X-axis shows sRNA abundances (RP10M) in corresponding sRNA libraries and the strand (W or C) to which they map. The Y-axis indicates the position (5’-terminal nucleotide position) of mapped sRNAs. B. Small RNAs mapped to the six gRNAs. The reads are from combined libraries of CaMV-inoculated *pCUT CP-1* and -2 at 3 dpi. The top panel shows an IGV screenshot for the collapsed view for gRNA1. The red box indicates the region that is enhanced in the six lower panels, as this region varies among gRNAs because the seed region of guide RNAs are within the 5’ terminal 22 to 23 nucleotides, while the remaining sequence is an invariant backbone. Gray shading at the top indicates coverage of sRNAs; red and blue colors indicate reads generated from “+” and “-” strands of the backbone. The lower six panels display small RNAs from the viral-specific seed regions of the six gRNAs; the gray dotted lines indicate the border of the seed region. Small RNAs were generated from the seed region of gRNA5; for gRNA6, of the few reads generated from gRNA6 seed region, the 3’ portion of nearly all corresponds to the non-seed backbone, meaning that these reads have little homology to viral RNAs. Small RNA sizes are indicated by colored dots as in Panel A, at the left end of each read in the lower six panels.

To further test whether gRNAs produce siRNAs, we mapped siRNA reads to these six gRNA sequences. Few siRNAs were generated from the 5’-terminal target site-specific regions of gRNAs; in contrast, abundant 21-nt siRNAs were derived from the antisense strand of a region close to the 3’ end of the gRNAs (Figure 2B). It is likely that these siRNAs are from dsRNAs and are dependent on the activity of RNA-dependent RNA polymerases and likely DCL4 (dicer-like 4) due to their 21-nt length. We performed siRNA target prediction against the CaMV RNA using gRNA-derived siRNAs, and these abundant siRNAs were not predicted to target the viral RNA, because of a lack of sequence complementarity. In addition, there were no siRNAs mapped to the 5’ spacer region of gRNA1-4. We observed only three 24-nt and one 22-nt siRNAs derived from the 5’ spacer region of the gRNA5 and one 21-nt siRNA mapped to that of gRNA6 (Fig. 2B). However, the abundance of these latter siRNAs was much lower than the sum of all siRNAs targeting CaMV genome, and therefore they seem unlikely to play a role in suppressing CMV RNAs. In summary, siRNA data confirmed that enhanced viral resistance is unlikely to be the result of siRNA-mediated silencing of viral RNAs. However, cellular RNAi could regulate the abundance of gRNAs by producing siRNAs concentrating at the 3’ end of gRNAs, in particular, when multiple gRNAs are present in a single cell.

We next tested the editing events at earlier and later stages during viral infection. For the earlier stage, we harvested and extracted DNA from CaMV-inoculated local leaves of *CUT CP* plants at three DPI. We amplified and cloned the *CP* coding sequences into a cloning vector. Among the 129 clones we tested, 56 of them had shorter *CP* fragments based on restriction enzyme digestion. Sequencing of 8 randomly picked clones (of the 56) showed large deletions in the CP coding sequence, ranging from 590 base pairs (bp) to 1351 bp (Fig. 3A). The large deletions were consistent with Cas9-mediated editing events and the expression of six gRNAs targeting the N-terminal 440 bp of the CP coding sequence (Fig. 3A). However, the deleted sequences, in most cases, extended beyond the 440 bp region and in several cases, beyond even the *CP* coding sequence. The 43% of clones that contained deletions is an underestimate of editing events because deletions caused by editing might have extended over the region covered by the pair of primers utilized. For the later infection stage, we used the systemically infected leaves of two *CUT CP* symptomatic plants. Among the 13 randomly sequenced *CP* gene clones, we recovered eight clones with wt *CP* sequences, and five mutants with short deletions or insertions consistent with Cas9-induced DNA breaks at the gRNA target sites in *CP* (Fig. 3B). Insertions were derived from duplications events in CaMV *CP* sequence. Many of the mutant CaMV encode CP protein predicted to be non-functional (Table S1). In these cases, we always recovered wt *CP* genomes as well, suggesting that mutant CaMV is likely packaged in virions via wt CP in *trans*. It should be noted that we did not identify any mutants with large deletions. It is likely that the large deletion was detrimental to CaMV survival and thus, such mutants were lost during viral infection.

**Figure 3.**
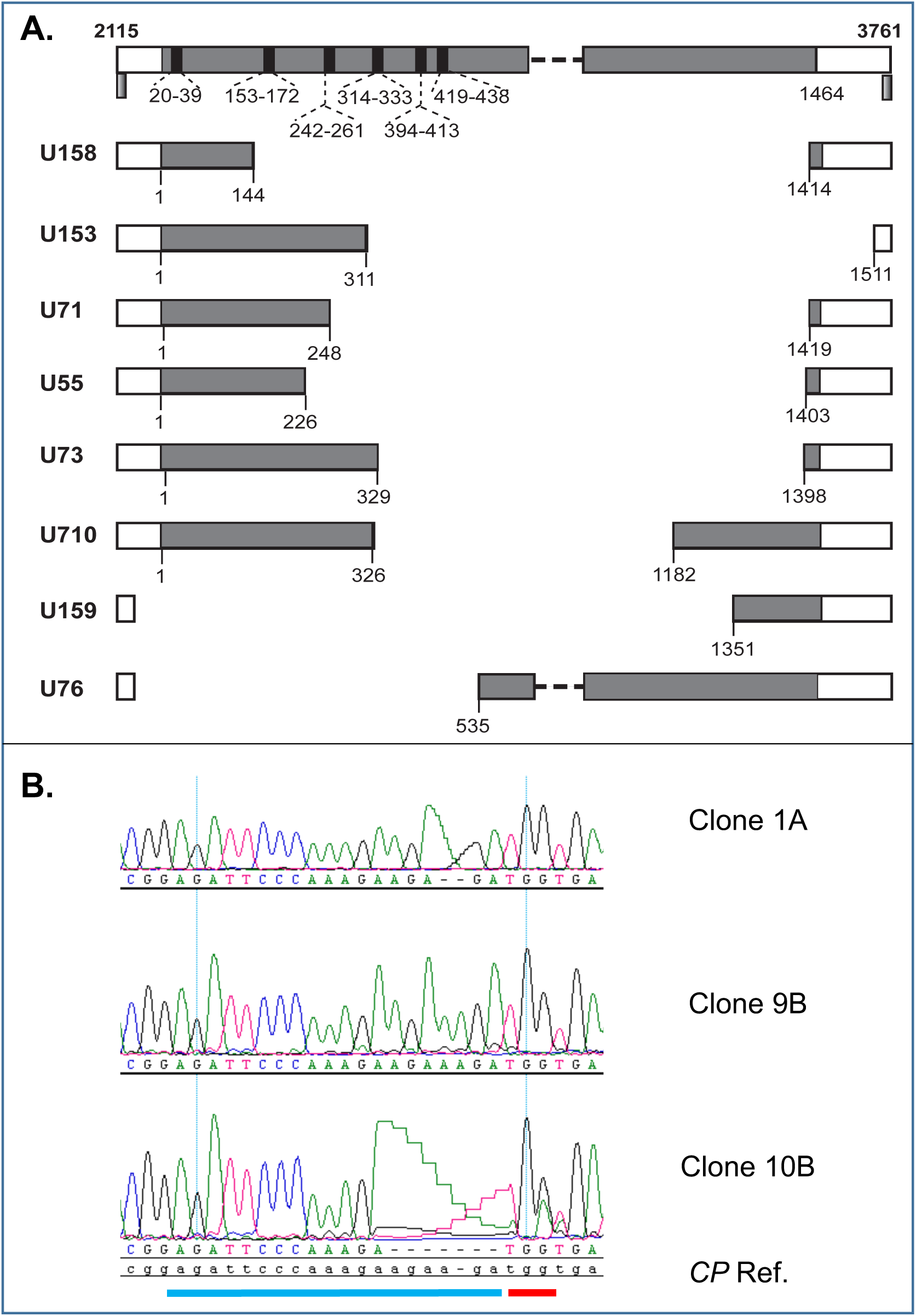
The editing events in the local and systemic leaves of *CUT CP* transgenic plants infected with CaMV. A. Large deletions identified in CaMV genome recovered from the inoculated leaves at 3 DPI. Numbers on top of the bar represent positions in CaMV genome. Numbers below bars show positions in the *CP* coding sequence, which is colored as grey. The two primers used to amplify the CaMV fragment of 2115-3761 encompassing the *CP* gene are identified as gradient bars. B. CaMV-induced Cas9 editing events from systemic tissues of *CUT CP* showing symptoms. Chromatographs from the sequenced CaMV *CP* gene cloned from *CUT CP* plants displaying symptoms. Blue bar, gRNA target sequence (G5), red bar (PAM site).

The complete resistance of the majority of *CUT CP* plants to CaMV infection suggests that Cas9-mediated viral targeting can be an effective strategy to control CaMV infection. Cas9-directed virus resistance has been achieved in several classes of human viruses, including reverse transcribing viruses HIV and Hepatitis B virus, as well as dsDNA viruses including human papillomavirus and Epstein-Barr virus (Price et al., 2016). Viral resistance to several plant geminiviruses conferred by CRISPR-Cas9 has also been reported (Ali et al., 2015; Baltes et al., 2015; Chaparro-Garcia et al., 2015; Ji et al., 2015), indicating the broad-range of applicability to confer resistance to viruses that viral dsDNA plays a critical role during their infections. However, virus mutants that escape CRISPR-Cas9-mediated resistance have been reported in several cases and in addition, it has been reported that different target sites may result in different levels of viral escape events. For instance, gRNAs targeting the coding sequences of plant geminiviruses led to frequent viral escapes, however, no viral escape was identified with gRNAs targeting the non-coding intergenic region, which is essential for replication (Ali et al., 2016). This is different from what has been previously reported regarding the Cas9-mediated resistance to HIV (Wang et al., 2016a; Wang et al., 2016b; Wang et al., 2016c). Escaped HIV mutants have been generated in Cas9-expressing cells and single gRNAs targeting either coding sequences or non-coding long terminal repeats (LTRs). However, gRNAs targeting highly conserved sequences, either coding or non-coding sequences, resulted in lower levels of escape mutants than those non-conserved sequences (Wang et al., 2016b; Wang et al., 2016c). Of note, the combinations of two single gRNAs, especially those far from each other, substantially delayed the resistance breakdown compared to single gRNAs. In particular, certain combinations of two gRNAs permanently blocked viral infection (Wang et al., 2016a). In the present work, we generated transgenic plants expressing both Cas9 and six gRNAs targeting CP sequences within a fragment of ~440 bp. We found large deletions within *CP* in the CaMV-inoculated leaves as early as three DPI. These deletions extended beyond the *CP* region containing all gRNA sequences and are not consistent with DNA repairs that are mediated by non-homologous end-joining. This could be due to multiple gRNAs concentrating at a short DNA fragment, introducing unexpected deletions. It is also possible that repairing the circular CaMV genome with multiple breaks generated by multiple gRNAs maybe different from the non-circular DNA (either as part of chromosome or linear DNA fragment). Large deletion caused by expressing two gRNAs has been reported during infection of geminivirus bean yellow dwarf virus. The majority of deletions occur between two gRNAs with intact sequences flanking gRNA-targeted sites, however, one of six clones had an extended deletion beyond the gRNA targeting sequence (Baltes et al., 2015). Although the rate of extended deletion is lower than that in our report, this points out the possibility that multiple gRNAs targeting circular DNA may result in unexpected deletions.

Our work also demonstrates that modified viruses are present in systemically infected leaf tissues, raising the possibility that Cas9-mediated viral targeting has the potential to create and allow the dissemination of mutant viruses from plants even with multiple gRNAs. The presence of duplication events suggests that Cas9 can promote recombination within viral genomes. This observation has implications for Cas9-mediated viral immunity strategies in plants. Targeting the *CP* of CaMV likely allows the escape of mutant viruses through the acquisition of wt CP function in *trans* from co-infecting viruses. Targeting of essential cis-acting regions of the viral genome might help prevent the escape of modified viruses based on previous geminivirus studies (Ali et al., 2016), in which Cas9 editing reduced viral accumulation and plant symptoms. While our current gRNA system allows up to 14 gRNAs in one vector (Peterson et al., 2016), a higher order multiplex strategy based on tRNA processing has been reported (Xie et al., 2015), enabling the targeting of multiple conserved cis- and trans-elements at multiple sites in various species. It is possible that arrays could be designed to target conserved sites among different viral strains or be combined to target multiple strains, or different viruses, making resistance broad spectrum as well as durable. The use of RNA-targeting Cas9 variants (Price et al., 2015) may additionally allow for the targeting of RNA viruses.

When controlling viruses using the CRISPR-Cas9 system, both Cas9 and gRNAs are consistently expressed in the cells. Recruiting Cas9 to viral DNAs depends on the presence and abundance of gRNAs. However, with the presence of folded dsRNA domains in gRNAs, siRNAs can be produced to contain the foreign RNAs. Based on deep-sequencing of small RNAs, we provided evidence that abundant 21-nt siRNAs were derived from the antisense strand of a region close to the 3’ end of the gRNAs (Figure 2B). The 3’ end of gRNAs are identical among all gRNAs. Our data suggest that the cellular RNAi pathway could affect the abundance of gRNAs and should be considered in the future application of the CRISPR-Cas9 system.

## Experimental procedure

### Cas9 target site selection and vector creation

Target sites in the CaMV *CP* gene (strain w260) were selected using standard Cas9 bioinformatic criteria (Table 1). Linear arrays of Arabidopsis *U6 promoter::gRNA* units were designed and subsequently synthesized by GeneArt/LifeTechnologies. Arrays were synthesized in groups of four, and combined by stacking using restriction enzyme cloning (Peterson et al., 2016). The arrays were cloned into the *pCUT3* binary vector, as were control *GLV* arrays containing 14 unique gRNAs (Peterson et al., 2016). As a control, the *CP* gRNA array was cloned into the parental *pCUT3* plasmid lacking *Cas9*.

### Plant growth and transgenic line selection

The constructs were transformed into an *Agrobacterium tumefaciens* recA-strain via triparental mating. Wild-type *Arabidopsis thaliana* Col-0 plants grown under constant light at 21°C were transformed via the floral dip method (Clough and Bent, 1998). T_1_ seeds derived from the transformed plants were collected and transgenic seedlings were selected on B5 media containing 100mg/L kanamycin. Kanamycin resistant seedlings were transferred to soil (Sunshine #1), and grown under 21 °C, 16-hour light/8- hour dark conditions. T1 plants of *CUT CP/No Cas*, T2 and T3 plants of two *CUT CP* transgenic lines (*CUT CP*-1 and -2) were used for CaMV infections.

### Viral Infection

Viral particles were isolated as described in (Schoelz et al., 1986) and infected into plants using mechanical inoculation. Two primary leaves were inoculated per plants and were marked. Newly emerged systemic leaves or inoculated leaves at the specified days post infection were harvested for DNA extraction.

### Analysis of CaMV DNA levels

The presence of CaMV was determined by PCR (Figs S1 & S2) from DNA isolated from systemic plant tissue in infected plants using primers specific to the CaMV *CP* or *RT* coding sequences (Table 2). As an amplification control, primers (VTXW934 & 935) for Arabidopsis *ACTIN2* gene were used for each DNA sample (Table 2). DNA was extracted by grinding leaf tissues in buffer TNES [0.2 M Tris-HCl (pH 7.5), 0.25 M NaCl, 0.025 M EDTA (pH 8.0), and 0.5% SDS], followed by phenol and chloroform extractions. To sequence the CaMV *CP* coding sequences, the PCR product was cloned to EcoRV-digested pBluescript KS plasmid (Stratagene, Agilent). For real-time PCR quantification (Fig 1), the primer pair VTXW1306 & 1307 was used to detect the *ACTIN2* gene and the primer pair VTXW1308 & 1309 was used to detect CaMV *CP*. DNA amplification was performed in the presence of iTaq Universal SYBR Green superMix (Biorad) using Applied Biosystems 7500 real time PCR system. Amplification conditions were: 95 °C for 2 min; 40 cycles of 15 seconds at 95°C and 60 seconds at 60 °C. Melting curve analysis was included to verify specificity of the amplification. Copy number of each gene was calculated according to standard curves, which were generated from a serial dilution of the plasmid pJT-ACT2 or pKS-CaMV CP. The ratios of CaMV CP vs. *ACTIN2* were calculated to demonstrate the relative accumulation of CaMV viruses.

**Table 2.**
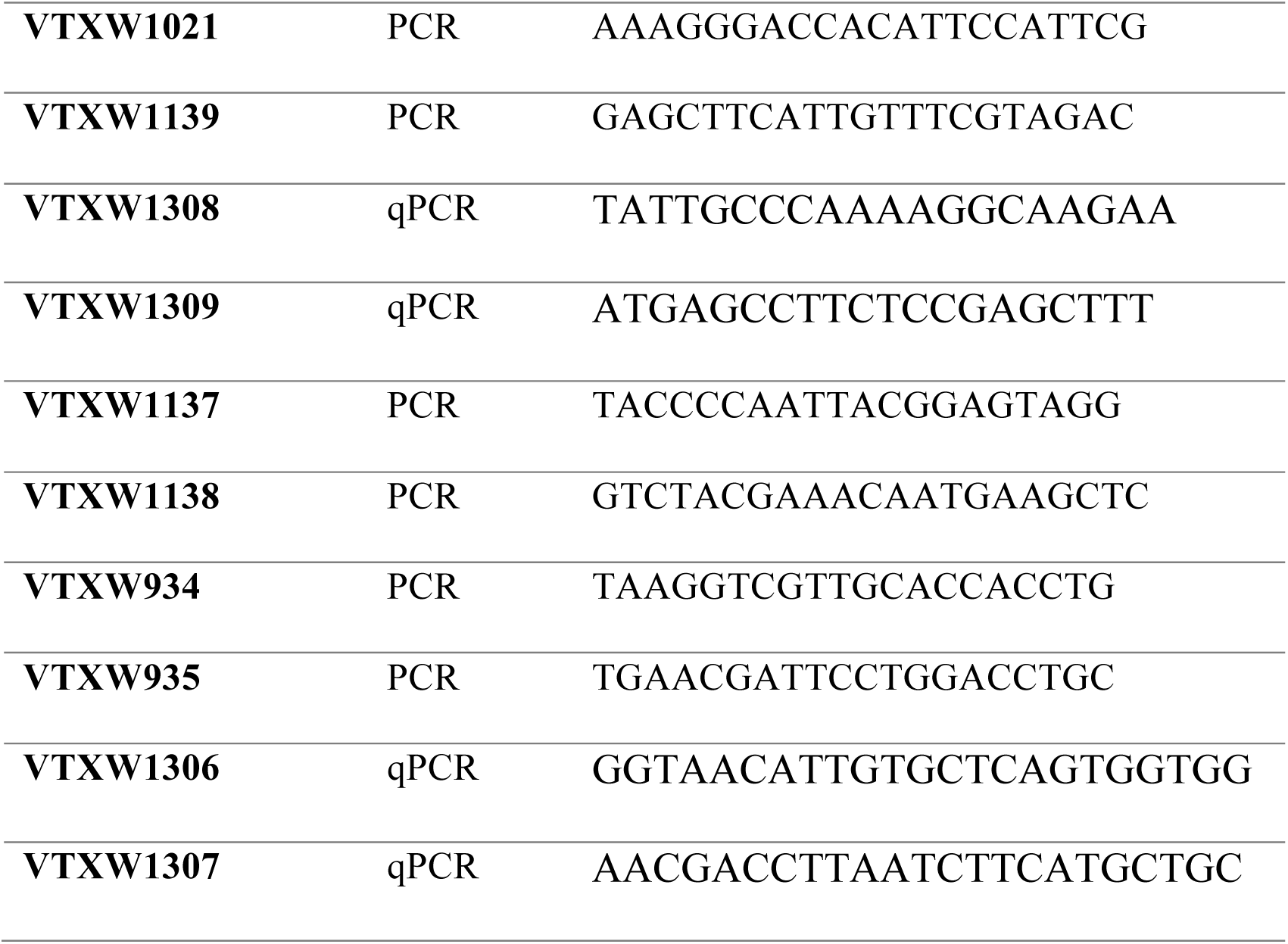
PCR primers for host *actin*, CaMV *CP* and *RT*.

### Small RNA library construction and analysis

Total RNA was isolated from Arabidopsis leaf samples using PureLink Plant RNA Reagent (Ambion). Eight sRNA libraries were prepared using TruSeq Small RNA Library Preparation Kits (Illumina). Eight small RNA libraries were constructed and were sequenced using an Illumina HiSeq 2500 at the Delaware Biotechnology Institute. Adapters in the raw sequencing data were trimmed. Small RNA reads between 18 nt and 30 nt were retained and were normalized to 10 million for comparisons among the libraries. Small RNA reads were mapped to the CaMV DNA sequence and gRNA sequences allowing zero mismatches using Bowtie (Langmead et al., 2009). Target prediction was performed for sRNAs derived from gRNAs in CaMV-infected samples against the CaMV DNA sequence using ‘psRNATarget’ (Dai and Zhao, 2011).

## Author contributions

H.L., J.L., and G.H. did CaMV infections and analysis, C.L.S. and B.A.P. made transgenic plants and analysis, C.L.S. and J.L. did mutation analysis, Q.F. did small RNA sequencing and analysis, B.C.M., Z.L.N. and X.W. conceived the concept and wrote the manuscript.

## Acknowledgements

We wish to thank Dr. James Schoelz (University of Missouri) for providing detailed protocols for virion preparation. We thank Dr. Guillaume Pilot for providing the *pJET ACTIN2* plasmid and Dr. Elizabeth Grabau for critical review of the manuscript. This work was supported by startup funds from Virginia Tech to both Z.L.N and X.W, startup funds from The University of North Carolina at Chapel Hill and National Science Foundation IOS awards: 1455607 and 1546837 to Z.L.N, and NSF IOS award 1257869 to B.C.M. Work in X. W.’s lab is also supported by the Virginia Agricultural Experiment Station, the Hatch Program of NIFA, USDA and NSF IOS award 1645749.

## Supplemental figures

**Figure S1 The gRNA sequences in relation to CaMV coat protein coding sequence (strain w260).** Targeting sequences of six gRNAs are in bold and PAM sites are underlined.

**Figure S2. Cas9-mediated viral immunity by targeting the CaMV *CP* coding sequence.** A. CaMV-infected plants at 26 DPI. B. The CaMV *CP* gene is detectable in Col-0, *CUT GLV* and *CUT CP* plants displaying symptoms, but not in symptomless *CUT CP* plants.

## References

Ali, Z., Abulfaraj, A., Idris, A., Ali, S., Tashkandi, M. and Mahfouz, M.M. (2015) CRISPR/Cas9-mediated viral interference in plants. Genome Biol 16, 238.

Ali, Z., Ali, S., Tashkandi, M., Zaidi, S.S. and Mahfouz, M.M. (2016) CRISPR/Cas9-Mediated Immunity to Geminiviruses: Differential Interference and Evasion. Sci Rep 6, 26912.

Anandalakshmi, R., Pruss, G.J., Ge, X., Marathe, R., Mallory, A.C., Smith, T.H. and Vance, V.B. (1998) A viral suppressor of gene silencing in plants. Proc Natl Acad Sci U S A 95, 13079–13084.

Baltes, N.J., Hummel, A.W., Konecna, E., Cegan, R., Bruns, A.N., Bisaro, D.M. and Voytas, A.F. (2015) Conferring resistance to geminiviruses with the CRISPR–Cas prokaryotic immune system. Nature Plants 1.

Cecchini, E., Al-Kaff, N.S., Bannister, A., Giannakou, M.E., McCallum, D.G., Maule, A.J., Milner, J.J. and Covey, S.N. (1998) Pathogenic interactions between variants of cauliflower mosaic virus and Arabidopsis thaliana. Journal of experimental botany 49, 731–737.

Chaparro-Garcia, A., Kamoun, S. and Nekrasov, V. (2015) Boosting plant immunity with CRISPR/Cas. Genome Biol 16, 254.

Chapdelaine, Y. and Hohn, T. (1998) The Cauliower Mosaic Virus Capsid Protein: Assembly and Nucleic Acid Binding In Vitro. Virus Genes 17, 139–150.

Clough, S.J. and Bent, A.F. (1998) Floral dip: a simplified method for Agrobacterium-mediated transformation of Arabidopsis thaliana. Plant J 16, 735–743.

Dai, X. and Zhao, P.X. (2011) psRNATarget: a plant small RNA target analysis server. Nucleic Acids Res 39, W155–159.

Ding, S.W. (2010) RNA-based antiviral immunity. Nat Rev Immunol 10, 632–644.

Erickson, F.L., Holzberg, S., Calderon-Urrea, A., Handley, V., Axtell, M., Corr, C. and Baker, B. (1999) The helicase domain of the TMV replicase proteins induces the N-mediated defence response in tobacco. Plant J 18, 67–75.

Fernandez, A., Hilson, P. and Beeckman, T. (2013) GOLVEN peptides as important regulatory signalling molecules of plant development. Journal of experimental botany 64, 5263–5268.

Incarbone, M. and Dunoyer, P. (2013) RNA silencing and its suppression: novel insights from in planta analyses. Trends Plant Sci 18, 382–392.

Ji, X., Zhang, H., Zhang, Y., Wang, Y. and Gao, C. (2015) Establishing a CRISPR–Cas-like immune system conferring DNA virus resistance in plants. Nature Plants 1.

Jinek, M., Chylinski, K., Fonfara, I., Hauer, M., Doudna, J.A. and Charpentier, E. (2012) A programmable dual-RNA-guided DNA endonuclease in adaptive bacterial immunity. Science 337, 816–821.

Kang, B.C., Yeam, I. and Jahn, M.M. (2005) Genetics of plant virus resistance. Annu Rev Phytopathol 43, 581–621.

Kasschau, K.D. and Carrington, J.C. (1998) A counterdefensive strategy of plant viruses: suppression of posttranscriptional gene silencing. Cell 95, 461–470.

Langmead, B., Trapnell, C., Pop, M. and Salzberg, S.L. (2009) Ultrafast and memory-efficient alignment of short DNA sequences to the human genome. Genome Biol 10, R25.

Makarova, K.S., Haft, D.H., Barrangou, R., Brouns, S.J., Charpentier, E., Horvath, P., Moineau, S., Mojica, F.J., Wolf, Y.I., Yakunin, A.F., van der Oost, J. and Koonin, E.V. (2011) Evolution and classification of the CRISPR-Cas systems. Nat Rev Microbiol 9, 467–477.

Maule, A.J., Caranta, C. and Boulton, M.I. (2007) Sources of natural resistance to plant viruses: status and prospects. Mol Plant Pathol 8, 223–231.

Nekrasov, V., Staskawicz, B., Weigel, D., Jones, J.D. and Kamoun, S. (2013) Targeted mutagenesis in the model plant Nicotiana benthamiana using Cas9 RNA-guided endonuclease. Nature biotechnology 31, 691–693.

Padgett, H., Watanabe, Y. and Beachy, R.N. (1997) Identification of the TMV replicase sequence that activates the N gene–mediated hypersensitive response. Molecular Plant-Micobe Interactions 10, 709–715.

Peterson, B.A., Haak, D.C., Nishimura, M.T., Teixeira, P.J., James, S.R., Dangl, J.L. and Nimchuk, Z.L. (2016) Genome-Wide Assessment of Efficiency and Specificity in CRISPR/Cas9 Mediated Multiple Site Targeting in Arabidopsis. PLoS One 11, e0162169.

Price, A.A., Grakoui, A. and Weiss, D.S. (2016) Harnessing the Prokaryotic Adaptive Immune System as a Eukaryotic Antiviral Defense. Trends Microbiol 24, 294–306.

Price, A.A., Sampson, T.R., Ratner, H.K., Grakoui, A. and Weiss, D.S. (2015) Cas9-mediated targeting of viral RNA in eukaryotic cells. Proceedings of the National Academy of Sciences of the United States of America 112, 6164–6169.

Sander, J.D. and Joung, J.K. (2014) CRISPR-Cas systems for editing, regulating and targeting genomes. Nat Biotechnol 32, 347–355.

Schoelz, J., Shepherd, R.J. and Daubert, S. (1986) Region VI of cauliflower mosaic virus encodes a host range determinant. Mol Cell Biol 6, 2632–2637.

Seo, J.K., Wu, J., Lii, Y., Li, Y. and Jin, H. (2013) Contribution of small RNA pathway components in plant immunity. Molecular plant-microbe interactions: MPMI 26, 617–625.

Wang, A. and Krishnaswamy, S. (2012) Eukaryotic translation initiation factor 4E-mediated recessive resistance to plant viruses and its utility in crop improvement. Mol Plant Pathol 13, 795–803.

Wang, G., Zhao, N., Berkhout, B. and Das, A.T. (2016a) A Combinatorial CRISPR-Cas9 Attack on HIV-1 DNA Extinguishes All Infectious Provirus in Infected T Cell Cultures. Cell Rep 17, 2819–2826.

Wang, G., Zhao, N., Berkhout, B. and Das, A.T. (2016b) CRISPR-Cas9 Can Inhibit HIV-1 Replication but NHEJ Repair Facilitates Virus Escape. Mol Ther 24, 522–526.

Wang, Z., Pan, Q., Gendron, P., Zhu, W., Guo, F., Cen, S., Wainberg, M.A. and Liang, C. (2016c) CRISPR/Cas9-Derived Mutations Both Inhibit HIV-1 Replication and Accelerate Viral Escape. Cell Rep 15, 481–489.

Xie, K., Minkenberg, B. and Yang, Y. (2015) Boosting CRISPR/Cas9 multiplex editing capability with the endogenous tRNA-processing system. Proc Natl Acad Sci U S A 112, 3570–3575.

